# Learning causal biological networks with the principle of Mendelian randomization

**DOI:** 10.1101/171348

**Authors:** Md. Bahadur Badsha, Audrey Qiuyan Fu

**Affiliations:** Department of Statistical Science, Institute for Bioinformatics and Evolutionary Studies, Center for Modeling Complex Interactions, University of Idaho, Moscow, ID, USA

## Abstract

Although large amounts of genomic data are available, it remains a challenge to reliably infer causal (i.e., regulatory) relationships among molecular phenotypes (such as gene expression), especially when many phenotypes are involved. We extend the interpretation of the Principle of Mendelian randomization (PMR) and present MRPC, a novel machine learning algorithm that incorporates the PMR in classical algorithms for learning causal graphs in computer science. MRPC learns a causal biological network efficiently and robustly from integrating genotype and molecular phenotype data, in which directed edges indicate causal directions. We demonstrate through simulation that MRPC outperforms existing general-purpose network inference methods and other PMR-based methods. We apply MRPC to distinguish direct and indirect targets among multiple genes associated with expression quantitative trait loci.

## Introduction

Whereas experiments (e.g., temporal transcription or protein expression assays, gene knockouts or knockdowns) have been conducted to understand the causal relationships among genes^1,2^, or between an expression quantitative trait loci (eQTL) and its direct and indirect target genes^3^, it remains a challenge to learn causality directly from genomic data. It is even harder to learn (i.e., infer) a causal network, which may represent which genes regulate which other genes. We address this problem in this paper. Correlation (or association) is often used as a proxy of a potentially causal relationship, but similar levels of correlation can arise from different causal mechanisms (Models 1-4 in **Fig. 1a**). For example, between two genes with correlated expression levels, it is plausible that one gene regulates the other gene (Models 1 and 2 in **Fig. 1a**); it is also plausible that they do not regulate each other directly, but both are regulated by a common genetic variant (Model 3 in **Fig. 1a**).

**Figure 1:**
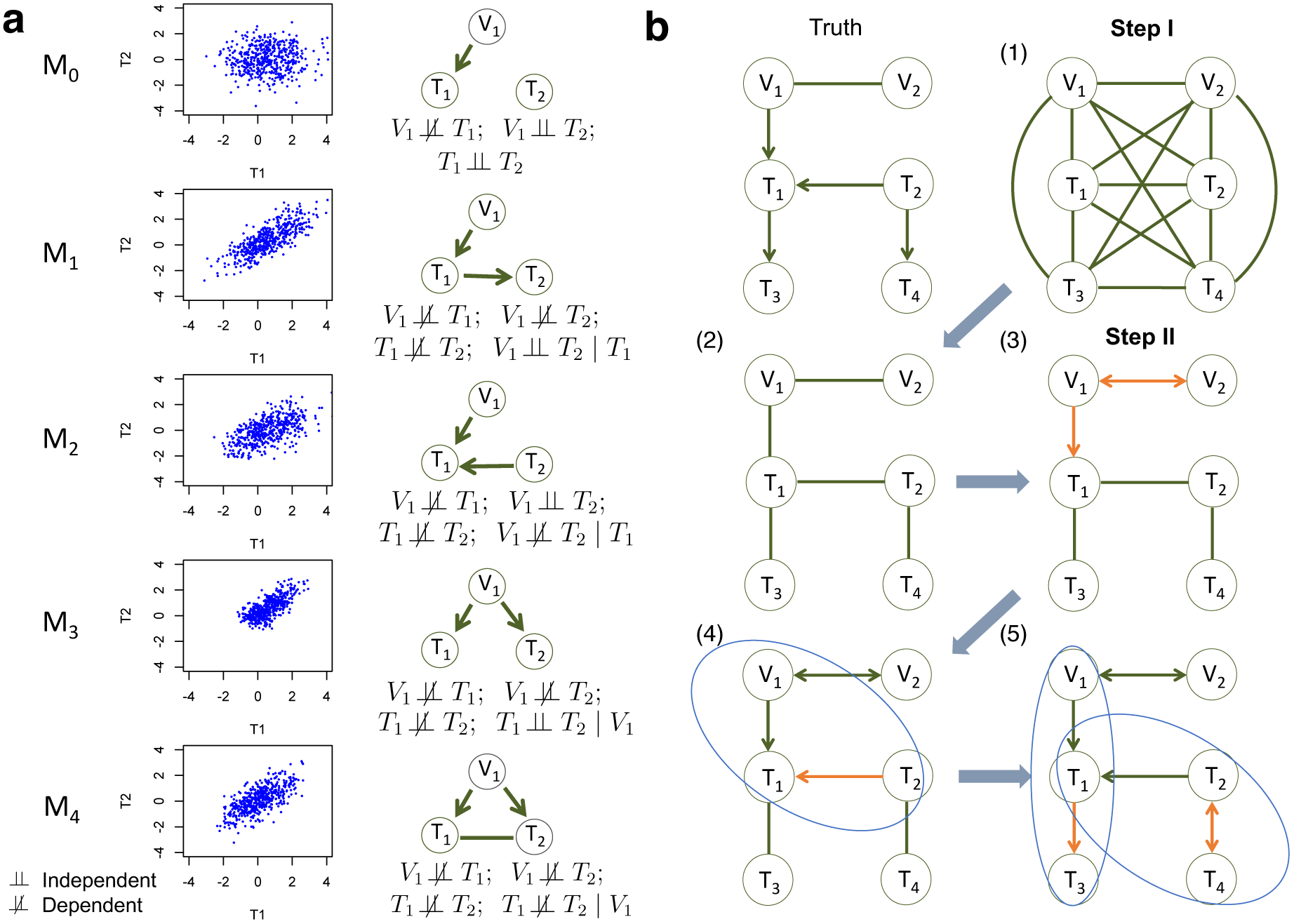
Five basic topologies under the principal of Mendelian randomization and the MRPC algorithm. **(a)** Each topology involves three nodes: a genetic variant (V_1_), and two molecular phenotypes (T_1_ and T_2_). Directed edges indicate direction of causality, and undirected edges indicate that the direction is undetermined (or equivalently, both directions are equally likely). For each topology (or model), a scatterplot between the two phenotypes is generated using simulated data, the topology is shown, and the marginal and conditional dependence relationships are given. M_0_ is the null model where T_1_ and T_2_ are marginally independent, and therefore the scatterplot does not show correlation. All the other models show scatterplots with similar levels of correlation. Our MRPC can distinguish the non-null models despite similar correlation. **(b)** The MRPC algorithm consists of two steps (see details in **Supplementary Figs 1 and 2**). In Step I, it starts with a fully connected graph shown in (1), and learns a graph skeleton shown in (2), whose edges are present in the final graph but are undirected. In Step II, it orients the edges in the skeleton in the following order: edges involving at least one genetic variants (3), edges in a v-structure (if v-structures exist) (4), and remaining edges, for which MRPC iteratively forms a triplet and checks which of the five basic models under the PMR is consistent with the triplet (5). If none of the basic models matches the triplet, the edge is left unoriented (shown as bidirected).

Correlation between the expression, or any molecular phenotype of two genes is symmetrical – we cannot infer which of the two genes is the regulator and which the target. However, if a genetic variant (e.g., a SNP) is significantly associated with the expression of one of the two genes, then we may assign a directed edge from the variant to the gene, as it is reasonable to assume that the genotype causes changes in the phenotype (expression), not the other way around. This additional, directed edge breaks the symmetry between the two genes, and makes it possible to infer the causal direction (e.g., compare Models 1 and 2 in **Fig. 1a**). This is the rationale behind the Principle of Mendelian Randomization (PMR). The randomization principle in experimental design (e.g., clinical trials) is critical in establishing causality: only when subjects are randomly assigned to different exposures is it possible to draw causal connections between exposure and outcome. As a randomization principle, the PMR assumes that the alleles of a genetic variant are randomly assigned to individuals in a population, analogous to a natural perturbation experiment and therefore achieving the goal of randomization^4^. The PMR has been widely used in epidemiology studies, where genetic variants are used as instrumental variables to facilitate the estimate of causal effect between a mediator (or exposure, such as gene expression) and an outcome (e.g., a disease phenotype^4^). It received increasing attention in genetics in recent years^5–17^

Large consortia, such as the GEUVADIS consortium^18^ and subsequently the GTEx consortium^19^, have established the widespread genetic variation (i.e. eQTLs) in human genome that may regulate gene expression, making PMR-based methods increasingly relevant and important for understanding interactions among genes. Furthermore, genome-wide association studies (GWASs) have identified a large number of genetic variants that are potentially causal to diseases^20^. Understanding the roles of these GWAS-significant variants is key to understanding the mechanisms underlying diseases. Interestingly, likely half of the GWAS-significant variants genetic variants are eQTLs^21^. As it becomes more common nowadays to collect gene expression data in disease studies^6,11^, studying eQTLs (which may also be GWAS-significant SNPs) and their associated genes provides a powerful approach for a deeper understanding of diseases.

However, existing methods adopting the PMR (e.g. the mediation-based methods^12,13^, and the MR methods^22^) are not directly applicable to inference of a causal network of gene expression. This is because these methods typically examine the graph of V_1_→T_1_→T_2_ (i.e., Model 1 in **Fig. 1a**), where V_1_ is the genetic variant, T_1_ may represent gene expression, and T_2_ a clinical trait. This graph, called the “causal model” by existing PMR-based methods, is sensible when T_2_ is a potential outcome of T_1_. However, when we examine relationships among gene expression or other molecular phenotypes, it is usually not known beforehand which of T_1_ and T_2_ is more likely to be the outcome of the other, and Model 1 alone does not have the flexibility of examining other possibilities. As a result, these methods are limited in the causal relationships they can recover. In this paper, we generalize the interpretation of the PMR to account for a variety of causal relationships.

On the other hand, in machine learning, a class of algorithms, such as those based on the classic PC algorithm^23–27^, have been developed in over a decade to efficiently learn causal graphs for a large number of nodes. These algorithms typically consist of two main steps (**Fig. 1b**): i) *inferring the graph skeleton* through a series of statistical independence tests. The graph skeleton is the same as the final graph except that the edges are undirected; and ii) *determining the direction of the edges* in the skeleton. Variants of the original PC algorithm have been developed to reduce the impact of the ordering of the nodes on inference (e.g., the R package pcalg^26,27^), or to reduce the number of statistical tests needed for inferring the skeleton (e.g., the R package bnlearn^24,25^).

Here we develop a new method, namely MRPC, which incorporates the PMR into PC algorithms and learns a causal graph where the nodes are genetic variants and molecular phenotypes (such as gene expression), and where the edges between nodes are undirected or directed, with the direction indicating causality. Crucially, by combining the PMR with machine learning, our method is efficient and accurate. Our extended interpretation of the PMR can be thought of as a way of introducing useful constraints in graph learning and effectively reducing the search space of topologies. We demonstrated the performance of MRPC on simulated and real data.

## Methods

### An extended interpretation of the Principle of Mendelian Randomization (PMR)

We extended the interpretation of the PMR to consider five causal relationships in a triplet of a genetic variant and two phenotypes, including the “causal model” (**Fig. 1a**). Under the assumption that genotype influences phenotype and not the other way around, these five models are mutually exclusive and encompass all possibilities, with Model 0 being the null model where the two phenotype nodes are not related, and the other four models being non-null models. As mentioned in the Introduction, Model 1 (V_1_→T_1_→T_2_) is typically referred to as the causal model under standard use of the PMR with T_1_ being the exposure (e.g., gene expression) and T_2_ being the outcome (e.g., clinical phenotype). cit^12,28^ and findr^13^, two existing PMR-based methods for example, both focus on testing Model 1.

However, Model 1 is limited. Among other possible causal relationships, Model 2 (V_1_→T_1_←T_2_) defines a v-structure where both edges point to the same node. This model is suitable when no genetic variant is available for T_2_ in the data. Model 3 (V_1_→T_1_ and V_1_→T_2_) captures the scenario where T_1_ and T_2_ are not directly related, but both regulated by V_1_. The current interpretation of the PMR in other methods typically rejects these two models in search of the “causal” model (Model 1). However, under our interpretation of the PMR, Models 2 and 3 describe alternative regulatory mechanisms between two genes, and therefore should also be allowed when constructing the network of molecular phenotypes. Model 4 (V_1_→T_1_; V_1_→T_2_; T_1_–T_2_) refers to the case where the two phenotypes T_1_ and T_2_ have additional dependence (represented by the undirected edge) on top of that induced by the sharing genetic variant. We consider undirected and bidirected edges to be equivalent for simplicity, in that an undirected edge can be thought of as representing two equally likely directions, namely M5 (V_1_→T_1_; V_1_→T_2_; T_1_→T_2_) and M6 (V_1_→T_1_; V_1_→T_2_; T_1_←T_2_). M5 and M6 are indistinguishable in terms of their dependence relationships (i.e., they are Markov equivalent^29^): all pairs of nodes can be marginally dependent and conditionally dependent given the remaining node. It is plausible that a hidden variable regulates both T_1_ and T_2_, although we currently do not consider hidden variables in our inference.

### MRPC, a novel causal network learning algorithm

Our method, namely MRPC, is a novel causal network inference method for genomic data (**Fig. 1b**; **Supplementary Figs. 1, 2)**. This method analyzes a data matrix with each row being an individual, and each column a genetic variant or a molecular phenotype. Our method also consists of the two main steps as described above. The first step of learning the graph skeleton is similar to that of other PC algorithms, but with an online control of the false discovery rate (FDR), which is explained in detail below. We incorporated the PMR in the second step of edge orientation (**Fig. 1b; Supplementary Fig. 2**), which involves three scenarios: i) MRPC first identify edges involving the genetic variants and orient these edges to point to the molecular phenotype; ii) MRPC then looks for three nodes with a potential v-structure (e.g., Model 2 in **Fig. 1a**, or among three molecular phenotypes, T_1_→T_2_←T_3_). MRPC conducts additional conditional independence tests if no such test has been performed in the first step; and iii) among the remaining edges, MRPC iteratively finds node triplets with only one undirected edge. It examines the results from the independence tests from the first step to identify which of the five basic topologies is consistent with the test results for this triplet. In MRPC, we use Fisher’s z transformation for Pearson correlation in all the marginal tests and for the partial correlation in all the conditional tests, consistent with the default method in pcalg (**Appendix A.1**). However, other parametric or nonparametric tests for marginal and conditional independence tests may be performed in place of Fisher’s z transformation test.

Existing network inference algorithms (such as those implemented in R packages pcalg and bnlearn) control the type I error rate for each individual statistical test, but not the family-wise error rate (FWER) or the FDR, as most methods controlling both the FWER and FDR require the knowledge of the total number of tests, which is not known in advance in graph learning. Lack of correction for multiple comparison often leads to too many false edges in the inferred graph, especially when the graph is large (see our simulation results below). We implemented in MRPC the LOND (Levels based on Number of Discoveries) method for controlling the FDR in an online manner^30^ (**Appendix A.2**). The LOND method estimates the expected FDR conditioned on the number of tests performed so far and the number of rejections from these tests.

Furthermore, genomic data may contain outliers^31^, which can greatly distort the inferred graph (see our simulation results below). Like pcalg, MRPC uses the correlation matrix, rather than the individual-feature matrix, as input. We implemented in MRPC a method for calculating the robust correlation matrix^31^ (**Appendix A.3**) in place of Pearson correlation to alleviate the impact of outliers if they are present.

## Results

### MRPC outperforms existing network inference algorithms and PMR-based methods on synthetic data in overall accuracy

We compared MRPC with two popular network inference algorithms: the pc method (implemented in pcalg) and the mmhc method (implemented in bnlearn), and three PMR-based methods, namely cit, findr and QPSO^32^. Except for QPSO, which is implemented in MATLAB, all the methods are implemented in R. We simulated data using linear models for the five basic topologies, three common topologies in biology^33,34^ (such as multi-parent, star, and layered graphs), as well as a complex topology with over 20 nodes (**Fig. 2**). We varied the sample size, as well as the signal strength through the coefficients in the linear models (**Appendix A.4**).

**Figure 2:**
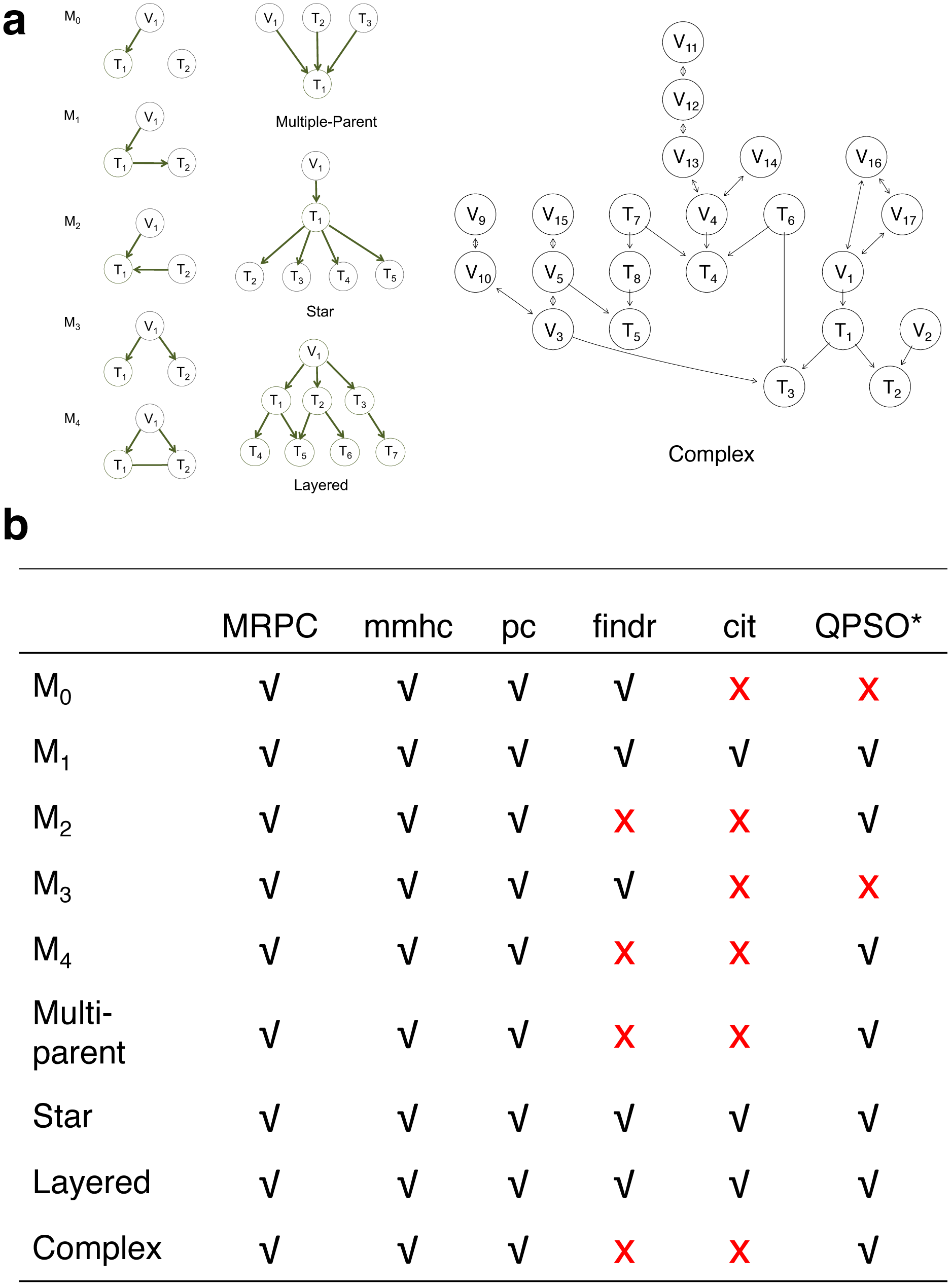
Comparison of MRPC with other methods on simulated data. **(a)** Topologies used to generate synthetic data (**Appendix A.4**). **(b)** Table summarizing graphs to which each method under comparison is applicable. *Note that QPSO does not learn the causal graph from scratch. Instead, it takes a graph skeleton as the input and seeks the optimal orientation of the edges in this undirected network. Edges involving genetic variants need to be already oriented in the skeleton. Therefore, QPSO does not identify M_0_ or M_3_.

For each topology, we generated 1000 data sets with different combinations of signal strength and sample size, and ran each method with their default parameters. Specifically, we ran MRPC with FDR=0.05, Pearson correlation (*β*= 0; see **Appendix A.2**) and the LOND method (*a* = 2; see **Appendix A.3**). We ran mmhc and pc with the type I error rate being the default value of 0.05. We explained the procedures for running other PMR-based methods in the next section.

We compared the recall and precision (**Appendix A.5**) across methods (**Fig. 3a, Supplementary Tables 1, 2, Supplementary Fig. 3)**. Recall (i.e., power, or sensitivity) measures how many edges from the true graph a method can recover, whereas precisions (i.e., 1-FDR) measures how many correct edges are recovered in the inferred graph. Across different topologies and parameter settings, MRPC has the highest median recall and precision, with both median recall and median precision above 80%. MRPC is followed by mmhc, QPSO, pc with two parameter settings, findr, with cit trailing far behind (**Fig. 3a**). MRPC recovers the true graph particularly well at moderate or stronger signal with a medium or larger sample size. For the complex topology, MRPC performs consistently better than pc and mmhc. This is still the case when the signal strength is heterogeneous across the complex topology (**Appendix A.6**; **Supplementary Fig. 4**). Examination of inferred graphs from different methods shows that pc is unable to determine edge directions or wrongly identifies v-structures when the true model contains none (**Fig. 3b**; **Supplementary Figs. 5, 6**). PMR-based methods, such as findr and cit, can infer too many or too few edges, whereas QPSO cannot identify the direction correctly. In the presence of outliers, MRPC with robust correlation as input substantially outperforms pc and mmhc (**Supplementary Fig. 7)**.

**Figure 3:**
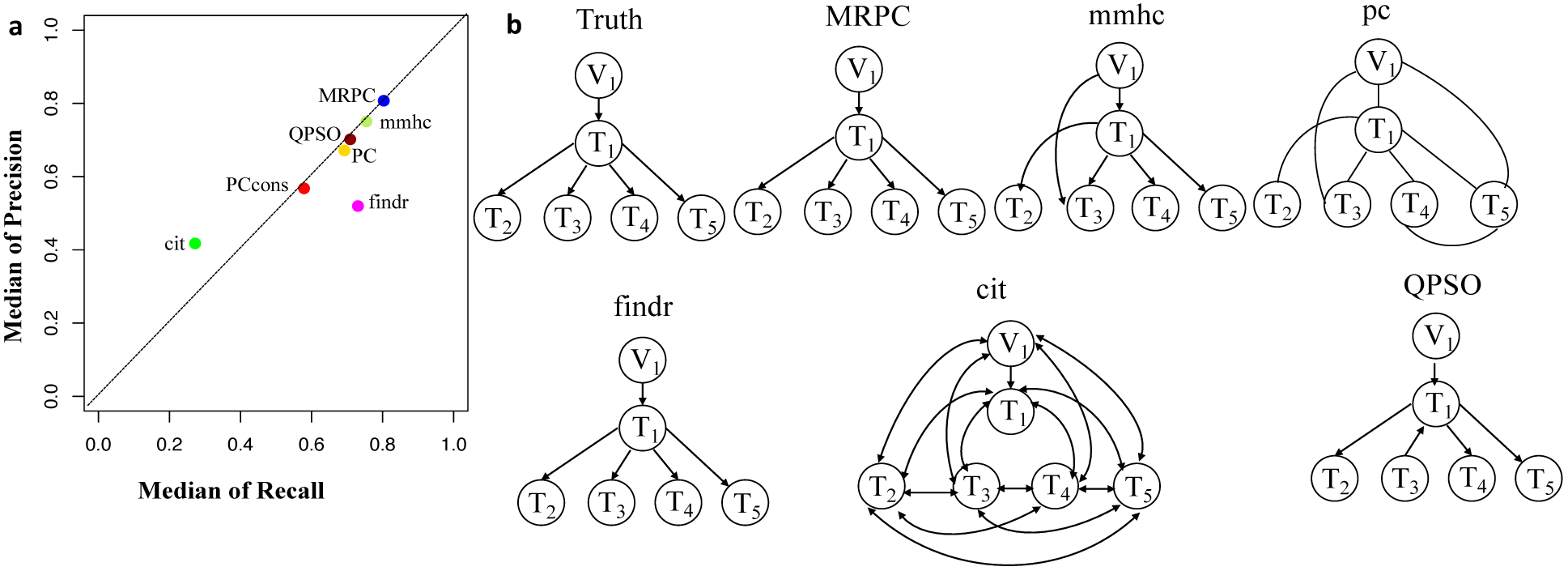
Results of method comparison on simulated data. **(a)** Median recall and precision over all parameter settings. For each of the topologies in **Fig. 2a**, 1000 datasets were generated for three different signal strengths (*γ*, which is the coefficient of parent nodes in the linear model; see Methods for simulation details) and four different sample sizes (*n*). Each of the six methods was applied where possible and the recall and precision were calculated for the inferred graph relative to the truth. The median of all the mean recall (or precision) is used as a metric of the overall performance of the method. We experimented with two settings of the pc function: the default (“PC”) and the conservative (“PCcons”). Since the default setting outperforms the conservative one, we use only the default setting in other analyses. Note that only 20 datasets were used for QPSO in each parameter setting due to long runtime. **(b)** An example of inferred graphs from all six methods on data simulated under a star model with a large sample size (*n* =1000) and strong signal (*γ* = 1.0).

### Existing PMR-based methods cannot deal with complex causal relationships

We examine the performance of PMR-based methods more closely in this section. Since cit and findr focus on Model 1, the topologies they can identify are limited to those that involve primarily Model 1, such as the star graph and the layered graph: the star graph consists of four M1s, and the layered graph five (**Fig. 2b**). For method comparison, we limited the true graphs to those that can be analyzed by findr or cit, specifically, M0, M1, M3, star and layered graphs for findr, and M1, star and layered graphs for cit (**Fig. 2b; Appendix A.7**).

Unlike MRPC, which is agnostic about which genes may be potential regulators and which potential targets, findr and cit are applied to ordered gene pairs iteratively, requiring specification of which of the two genes is the potential regulator and which the target. For example, to test whether the data are simulated under M1, then findr and cit will be performed twice, on (V_1_, T_1_, T_2_) and then on (V_1_, T_2_, T_1_). The number of ordered gene pairs is 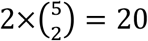 for the star graph and 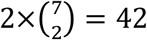 for the layered graph. We applied Bonferroni correction with a familywise type I error rate of 0.05. Take again the star model with a sample size of 1000 for example, where we varied the signal strengths in simulation. Although Bonferroni correction is already a conservative method for multiple testing, findr still sometimes infers more edges than there are (summarized by the lower precision in **Fig 3a**), whereas cit may infer a very dense graph or no edges at all (summarized by low recall and low precision in **Fig. 3a**; also see examples in **Fig. 3b** and **Supplementary Figs. 5, 6**). Even when the graph skeleton is known, findr and cit still did not outperform MRPC in nearly all cases (**Supplementary Figs. 8, 9**).

Similar to MRPC, QPSO also has connections to PC-like algorithms. However, QPSO does not infer a graph skeleton. Instead, it requires a graph skeleton as the input and seeks the optimal orientation of the edges, its performance therefore depending heavily on how well the skeleton is inferred. Whereas the authors of QPSO used pc to generate the skeleton, we used MRPC to generate the input, having observed the unsatisfactory performance of pc. With a more accurate skeleton, QPSO is still lacking both in recall and in precision **(Fig. 3a)**. Additionally, QPSO is at least an order of magnitude slower than other methods (**Supplementary Table 3**). We therefore calculated recall and precision only for 20 (instead of 1000) data sets in simulation for QPSO.

### Application of MRPC to distinguishing direct and indirect targets of eQTLs

We are interested in identifying true targets when a single SNP is statistically associated with the expression of multiple genes. Multiple genes are potential targets often because these genes are physically close to one another on the genome, and the eQTL analysis usually examines the association between one SNP-gene pair at a time, ignoring dependence among genes. Indeed, among eQTLs identified from the GEUVADIS data^18^ (i.e., gene expression measured in lymphoblastoid cell lines, or LCLs, of a subset of individuals genotyped in the 1000 Genomes Project), 62 eQTLs discovered under the most stringent criteria have more than one associated gene (**Appendix A.8**). We applied MRPC to each of these eQTLs and their associated genes in the 373 Europeans, and identified 11 types of topologies (**Fig. 4; Supplementary Table 4;** also see comparison with mmhc and pc for some of the eQTL-gene sets in **Supplementary Fig. 10)**. Three of these 11 types are Models 1, 3 and 4 shown in **Fig. 1a**. Seven other topologies are identified for eight eQTLs each with three associated genes (**Supplementary Table 4)**.

Although the multiple associated genes of the same eQTL are physically near one another, our method showed promise in teasing apart the different dependence (or regulatory relationships) among these genes. For example, the SNP rs479844 (chr11:65,784,486; GRCh38), one of the 62 eQTLs, turns out to be significant in at least three GWASs for atopic march and more specifically, atopic dermatitis (p values ranging from 10^-10^ to 10^-18^)^20, 35–37^. This SNP has been linked with two genes, AP5B1 (chr11:65,775,893-65,780,802) and OVOL1 (chr11:65,787,022-65,797,219), in these GWASs, but it is unclear which is the real target. Our MRPC infers Model 1 for the triplet: rs479844→OVOL1→AP5B1 (**Fig. 4a**), which suggests that OVOL1 is more likely to be the direct target, and AP5B1 the indirect one. Meanwhile, for HLA-DQA1 (chr6:32,637,403-32,654,846) and HLA-DQB1 (chr6:32,659,464-32,666,689), both genes are associated with the SNP rs9274660 and located in the major histocompatibility (MHC) region of high linkage disequilibrium. As expected, MRPC infers an undirected edge between the two genes, as the information on the two genes is highly symmetric in the genotype and gene expression data. By contrast, mmhc and pc often misspecify edges or their directions (**Supplementary Fig. 10**). We focused on the European sample in this analysis, as the sample size of the Africans is small (89). However, we managed to replicate part of the topologies for the few eQTLs discovered in both populations (**Appendix A.8**).

**Figure 4:**
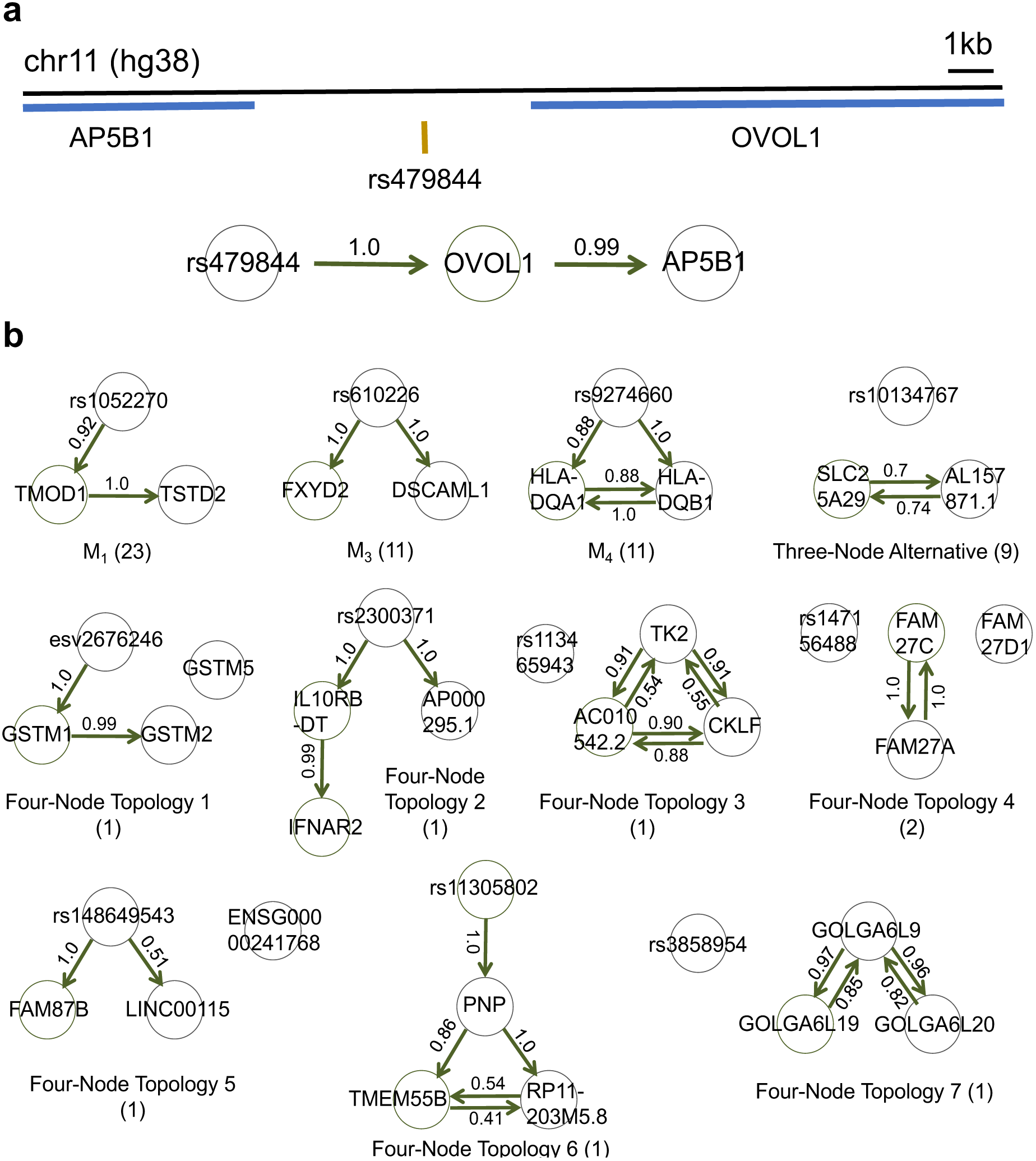
MRPC distinguishes direct and indirect target genes of eQTLs in the GEUVADIS data for the European cohort. **(a)** rs479844 is a GWAS significant SNP for atopic march in the GWAS Catalog, and an eQTL identified in GEUVADIS for two genes. (**b**) MRPC learns 11 distinct topologies among associated genes for eQTLs. Numbers on edges are probabilities of the corresponding directed edge being present in a bootstrap sample of 200. The number in parentheses under each topology is the number of eQTL-gene sets with the corresponding inferred topology.

Since the GTEx consortium^19^ contains data also from LCLs, we next examined whether the causal relationships inferred from the GEUVADIS data may be replicated in the LCL samples from GTEx (**Appendix A.9**). The sample size of 117 is much smaller in GTEx, though, which reduces the expected number of causal relationships to be replicated. We therefore focus on eQTL-gene sets that were inferred to have an M1 model in GEUVADIS by MRPC. We ran MRPC, findr and cit on the 16 eQTL-gene sets with an M1 model that have the genotype and gene expression data in both GEUVADIS and GTEx LCL samples. findr replicated 9 sets, MRPC 8 and cit only 1 (**Supplementary Table 5**). This result is consistent with simulation results (**Fig. 3a**): whether the graph skeleton is known or not, MRPC and findr have similar performance on M1 across different sample sizes and signal strengths, both much better than cit. In particular, we replicated the relationship rs479844→OVOL1→AP5B1 with both MRPC and findr in the GTEx LCL samples.

## Discussion

In summary, we have developed MRPC to infer causal networks, which can be an NP-complete problem (**Appendix A.10**). Our MRPC method examines a variety of causal relationships implied by the PMR, and takes advantage of the development of machine learning algorithms for causal graph inference. MPRC integrates genotypes with molecular phenotypes, and can efficiently and accurately learn causal networks. Our method is flexible as it requires only the genotype data (SNPs or other types of variants; see **Appendix A.11**) and the molecular phenotype measurements (gene expression, or other functional data, such as exon expression, RNA editing, DNA methylation, etc.), and can be applied to a wide range of causal inference problems. Our method is also nonparametric in that no explicit distributions are assumed for the underlying graph. MRPC uses individual-level genomic data to learn plausible biological mechanisms from combining genotype and molecular phenotypes.

The key improvements in MRPC over existing methods are i) implementation of the online FDR control method (the LOND method), which helps reduce false positives. As our simulation demonstrated, false positive edges are a severe problem in other methods, whether they are based on the PMR or not; and ii) accounting for all possible causal relationships a triplet with a genetic variant can have under the PMR. This extended interpretation of the PMR allows MRPC to go beyond the typical “causal model” examined by other PMR-based methods and can deal with networks of realistic causal relationships. Computationally, incorporation of the PMR puts constraints on the space of possible graphs and allows for efficient search of graphs consistent with the data (**Appendix A.12**).

Here, we demonstrated the outstanding performance of MRPC on small to moderately-sized graphs. Additional work is needed to extend the ability of MRPC to larger graphs while retaining inference accuracy. Indeed, apart from mmhc and pc, other existing methods for inferring large causal graphs also tend to have high false positive rates: for example, the TRANSWESD method developed for the DREAM5 Systems Genetics Challenge A (a network of 1000 SNPs and 1000 genes with directed and undirected edges) showed better performance than other participating method for this challenge. However, even TRANSWESD has an actual FDR as high as 64% at a large sample size of 999^48^, suggesting that much work is still needed to accurately infer a large causal graph.

Our current model behind MRPC also does not account for additional noise in measurements. By calculating correlation and performing statistical tests based on correlation, we account for variation in gene expression, which contains both (true) stochasticity in expression and measurement error. For future development, we will consider additional measurement errors that lead to systematic bias in the data^22^.

Like most causal graph learning methods, a key assumption behind MRPC is that there are no hidden or confounding nodes that are connected to the observed nodes in the graph. As the next step, we are working on extensions of MRPC to account for confounding variables. In our application here, the effect of confounding variable is alleviated as we focus on genes that have been identified to be strongly associated with the eQTL.

## APPENDIX A

### A.1 Conditional independence tests based on partial correlations

We use the same method (and R functions) as that used in the R package pcalg^26^ for conducting conditional independence tests based on partial correlations. Consider testing conditional independence between variables *x* and *y* conditioned on a set of variables *S*. From the correlation matrix, one may estimate the partial correlations using an iterative approach^26^. Then application of Fisher’s z transformation gives the test statistic

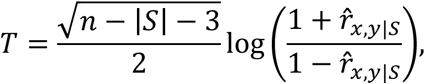

which follows N(0,1) under the null hypothesis of conditional independence^49^. In the expression above, *r*̂_*x,y*|*s*_is the estimated partial correlation, *n* the sample size, and |*S*| the number of variables in the set *S*.

### A.2 Sequential FDR control

We implemented the LOND algorithm that control FDR in an online manner, as we did not know the number of tests beforehand in learning the causal graph. Specifically, consider a sequence of null hypotheses (marginal or conditional independence between two molecular phenotypes) *H*(*m*)=*H*_1_,*H*_2_,*H*_3_,…,*H_m_*, with corresponding *p*-values p(m)=*p*_1_,*p*_2_,*p*_3_,…,*p*_m_. The LOND algorithm aims to determine a sequence of significance level *a_i_*, such that the decision for the *i*th test is

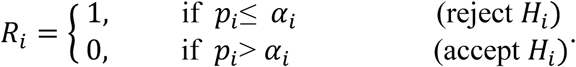

The number of rejections over *m* tests is then

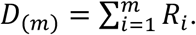

For the overall FDR to be *δ*, the significance level *α_i_* is set to be

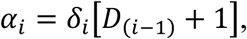

where the FDR for the *i*th test is

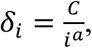

such that

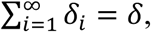

for integer *a*>1 and a constant *c*. We choose a nonnegative sequence *δ_i_*, such that 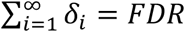. The default value for *a* is set to 2 in MRPC. At an FDR of 0.05 and *a* = 2, we have

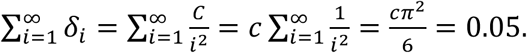

Then

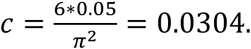

Values of *δ_i_* and *a_i_* for the first 18 tests of analysis of a simulated data set are listed in an example given in **Supplementary Table 6.** The larger *a* is, the more conservative the LOND method, which means that fewer rejections will be made. We therefore used *a* = 2 throughout simulation and real data analyses. Simulation results in the Results section show that this choice of *a* works reasonably well for small and moderately-sized networks.

### A.3 Calculation of robust correlation

We implemented the method in Badsha et al.^31^ to calculate the robust correlation matrix as the input to the MRPC inference. Specifically, for data that are approximately normal (usually after preprocessing of the data), we calculated iteratively the robust mean vector ***μ*** and the robust covariance matrix ***v*** until convergence. At the *t*+1 st iteration,

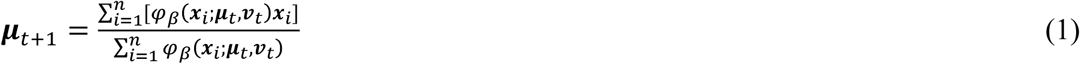

and

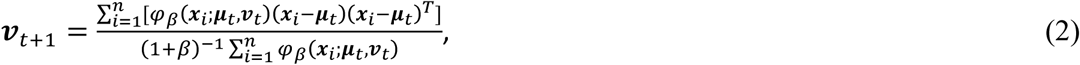

where,

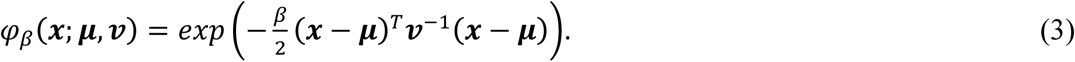

In the equations above, ***x***_*i*_ is the vector of gene expression in the *i*th sample, *n* the sample size, and *β* the tuning parameter. Equation (3) downweighs the outliers through *β*, which takes values in [0,1]. Larger *β* leads to smaller weights on the outliers. When *β* = 0, equation (2) is similar to the standard definition of the variance, except that the scalar is 1/*n*, whereas the unbiased estimator of the variance has a scalar of 1/(*n*-1). When the data matrix contains missing values, we perform imputation using the R package mice^50^. Alternatively, one may impute the data using other appropriate methods, and calculate the correlation matrix as the input for MRPC.

When analyzing simulated data with no outliers, we set *β*= 0, which is close to Pearson correlation. We set *β*= 0.005 if outliers were included in simulation. On real data, we would usually perform two analyses with *β* = 0 and *β* = 0.005. These two values did not lead to different results in most cases. See details in **Appendix A.8**, which refers to **Supplementary Figs. 11-13**.

### A.4 Generating simulated data

We generated synthetic data for a variety of graphs, which fall into three categories depending on the complexity (**Fig. 2a**): i) basic topologies of a triplet; ii) topologies common in biological networks: star (i.e., one molecular phenotype has multiple targets); multi-parent (i.e., one molecular phenotype has multiple regulators apart from the genetic variants); and layered; and iii) a complex topology.

In each topology, we simulated the data first for the nodes without parents, and then for other nodes. Genetic variants are nodes without parents, and we assume them to be biallelic SNPs with three genotypes 0, 1, and 2. Denote the minor allele frequency by *q* and assume Hardy-Weinberg equilibrium. Then the genotype of the *i*th variant, *V_i_*, follows a multinomial distribution:

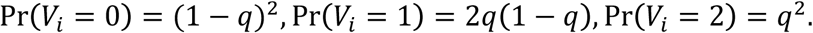

Denote the *j*th molecular phenotype by *T_j_* and the set of its parent nodes by P, which may be empty, or may include variant nodes or nodes of other molecular phenotypes. We assume that the molecular phenotype *T_j_* follows a normal distribution

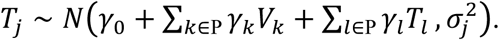

The variance may be different for different nodes. For simplicity, we use the same value for all the nodes.

We treat undirected edges as bidirected edges and interpret such an edge as an average of the two directions with equal weights. For example, for the undirected edge in Model 4 in **Fig. 1a**, we generate data for T_1_→T_2_:

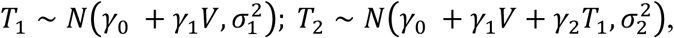

and separately for T1←T2:

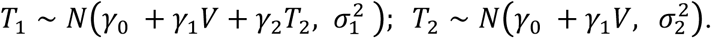

We then randomly choose a pair of values with 50:50 probability for each sample.

For simplicity in simulation, we set *γ*_0_ = 0 and all the other *γ* ′s to take the same value, which reflects the strength of the association signal. We considered three values for the slopes: 0.2 (weak signal), 0.5 (moderate signal), and 1.0 (strong signal). We also varied the sample size: 50 (very small), 200 (small), 500 (medium), and 1000 (large). Thus, we considered twelve combinations of signal strength and sample size (**Supplementary Tables 1, 2**).

Under each combination, we generated 1000 data sets for each topology. For each data set, we shuffled the columns corresponding to gene expression to generate one data set with those columns reordered; if an inference method is sensitive to the ordering of the columns, the inferred graph would have a large variance across data sets. We then applied each method to a data set with permuted columns. To summarize the results, we computed the mean and standard deviation of recall and precision (**Appendix A.5**) across 1000 data sets for each method. The median of recall and precision of each method across all topologies and all parameter settings are displayed in **Fig. 3a**. We also summarize the median standard deviation of recall and precision in **Supplementary Fig. 3**. Note that the standard deviation in recall and precision reflects variation due to both different data sets and different node orderings. Except for QPSO, the methods under comparison do not differ much in variation. QPSO had a larger variation because only 20 data sets were used for assessing the performance (due to long runtime).

### A.5 Recall and precision

Under the standard definition:

Recall = (# edges correctly identified in inferred graph) / (# edges in true graph);

Precision = (# edges correctly identified in inferred graph) / (# edges in inferred graph).

However, we consider it more important to be able to identify the presence of an edge than to also get the direct correct. Therefore, we assign 1 to an edge with the correct direction and 0.5 to an edge with the wrong direction or no direction. For example, when the true graph is V→T_1_→T_2_ with 2 true edges, and the inferred graphs are i) V→T_1_→T_2_, and V→T_2_; ii) V→T_1_–T_2_; and iii) V→T_1_←T_2_, the number of correctly identified edges is then 2, 1.5 and 1.5, respectively. Recall is calculated to be 2/2=100%, 1.5/2=75%, and 1.5/2=75%, respectively, whereas precision is 2/3=67%, 1.5/2=75%, and 1.5/2=75%, respectively.

When analyzing the complex topology in simulation, which involves correlated genetic variants, we ignored the edges among genetic variants in the calculation of recall and precision, since mmhc and pc are not designed to infer the relationships among genetic variants correctly. findr and cit are not applicable to this topology, and QPSO requires the graph skeleton as the input, with the graph skeleton already specifying the relationship among genetic variants and between variants and their associated genes.

### A.6 Simulation under the complex topology with heterogeneous signal strengths

The simulation strategy described above assumes the same signal strength (value of *γ*, the coefficient of the parent node) across the network, which allows us to examine the performance of the methods in simple and well-controlled settings. For the complex strategy, we further allowed the values of *γ* to vary when generating data for each node. Each *γ* has equal probability of taking on one of three values: 0.2, 0.5 and 1.0. Similar to the procedure described above, we also generated 1000 data sets with this strategy, applied relevant methods, and computed recall and precision.

### A.7 Application of findr and cit

Unlike mmhc and pc that learn the graph skeleton first and orient the edges next, findr and cit test for directed edges in a single step for a triplet of nodes (the genetic variant and two gene expression nodes). This means that in order to learn the topology, we needed to examine all possible gene pairs (e.g., T_1_ and T_2_; and T_2_ and T_1_) and then apply findr or cit to the triplet of each of the gene pairs and the genetic variant. Based on the hypothesis testing result from findr or cit, if there was evidence for a directed edge between two nodes, we added 1 to the current value in the adjacency matrix for those two nodes. Otherwise we left the value unchanged. After examining all gene pairs, we converted all positive values in the adjacency matrix to 1 to represent a directed edge. This way, no edges inferred would be eliminated in later tests. We then calculated recall and precision using the inferred adjacency matrix and that of the true graph, and averaged the rates across simulated data sets.

Although findr aims to compute a causality probability for a triplet, its current implementation for this calculation cannot be applied to small graphs, or cases where multiple genes share the same eQTL and where some of the genes do not have eQTLs. We therefore used the function findr.pijs_gassist_pv() from the R package findr to conduct five hypothesis tests (the p values from these five tests are then converted to a causality probability) for each ordered gene pair with the genetic variant. Consider a triplet V_1_, T_1_ and T_2_. The null (H_0_) and alternative (H_a_) hypotheses of these five tests are:

Test #1: H_0_: V_1_ and T_1_ independent; H_a_: V_1_→T_1_;
Test #2: H_0_: V_1_ and T_2_ independent; H_a_: V_1_→T_2_;
Test #3: H_0_ (M1): V_1_→T_1_→T_2_; H_a_: V_1_→T_1_, V_1_→T_2_, T_1_→T_2_;
Test #4: H_0_ (M0): V_1_→T_1_, both independent of T_2_; H_a_: V_1_→T_1_, V_1_→T_2_, T_1_→T_2_;
Test #5: H_0_ (M3): V_1_→T_1_, V_1_→T_2_; H_a_: V_1_→T_1_, V_1_→T_2_, T_1_→T_2_.

We extract the p values (i.e., *p_i_*, *i* = 1,…, 5) for the five tests. The data supports M0, if *p*_1_ is less than, and *p*_2_ and *p*_4_ greater than a certain threshold. The data supports M1, if *p*_1_ is less than, and *p*_3_ greater than a certain threshold. The data supports M3, if *p*_1_ and *p*_2_ are less than, and *p*_5_ greater than a certain threshold. We determine the p value threshold with Bonferroni correction, dividing the unadjusted p value 0.05 by 5*m*, where *m* is the total number of genes pairs, because each findr test contains five tests.

cit generates an omnibus p value for testing whether the triplet follows M1. We used the function cit.cp() from the R package cit for calculation of the omnibus p value. Similarly, we determine the p value threshold also with Bonferroni correction (unadjusted p value 0.05 divided by the total number of genes pairs).

When the graph skeleton is known, the number of tests is reduced to one on simple models (M0, M1 and M3), and to four in the star graph and to five in the layered graph (**Supplementary Figs. 8, 9**). In other words, potential regulators and targets are known to findr and cit. For MRPC we continued to assume that the skeleton was unknown. With known skeletons, both findr and cit performed similarly to, and in almost all the cases not better than MRPC. The performance of cit can still be much worse than the other two when the signal strength is low or the sample size is small.

### A.8 Analysis of the GEUVADIS data

The GEUVADIS project (http://www.ebi.ac.uk/Tools/geuvadis-das/) performed RNA-seq (gene expression) on 373 Europeans and 89 Africans from the 1000 Genomes Project. The GEUVADIS project combined the gene expression data with the genotype data, and identified eQTLs across the human genome. Among the most stringent set of eQTLs, 62 have more than one target gene. We extracted the genotypes of these eQTLs and the expression of the target genes in the 373 Europeans, and applied MRPC to each eQTL with its target genes.

The SNP rs479844, which has GWAS significance, is identified in the European sample to be the best eQTL for genes OVOL1 and AP5B1. However, this SNP is not identified to be the eQTL of any gene in the African sample. No eQTLs are reported for these two genes in the African sample. When we further examined the correlation matrices (**Supplementary Figure 11**) between the SNP genotype and expression of the two genes from the two samples, they have qualitative differences: whereas the eQTL has a much stronger correlation with OVOL1 than with AP5B1 in Europeans, it is the reverse in Africans. However, these differences are likely due to the small sample size of the African sample. We therefore do not seek to replicate with the African sample the topology we identified in the European sample.

Also because of the small sample size of the African sample, eQTLs and genes identified to have eQTLs are very different in the two populations. In order to examine whether it is possible at all to replicate the causal network inference from the European sample, we focused on the five top eQTLs identified in both samples: namely, esv2658282, esv2676246, rs11305802, rs230326, and rs7663027. The pairwise correlation matrices (**Supplementary Figure 12**) for each eQTL in the two samples are largely similar. However, due to the difference in the sample size, the topology inferred from the African sample is usually part (**Supplementary Figure 13**) of that from the European sample.

### A.9 Analysis of the GTEx data

The GTEx consortium has profiled genotypes and gene expression levels in 53 tissues across 714 donors (Release V7, dbGaP Accession phs000424.v7.p2; https://www.gtexportal.org/home/). We extracted the gene expression data of the LCLs, and the genotype data of the eQTLs used in the GEUVADIS analysis. Since GTEx uses chromosome locations to identify genetic variants, we extracted the coordinates of the GEUVADIS eQTLs in Ensemble (GRCh 37; https://grch37.ensembl.org/index.html) using the rs IDs. Not all GEUVADIS eQTLs can be found in the GTEx samples. Among eQTLs that can be found in the GTEx samples, not all their associated genes have expression measurements. In the end, we found 40 eQTL-gene sets with data available in both GEUVADIS and GTEx LCLs (**Supplementary Table 4)**. For each of these sets, we ran MRPC with an FDR of 0.05, and summarized the results in **Supplementary Table 4**. For those sets that were inferred to have an M1 model by MRPC in GEUVADIS, we also ran function findr.pijs_gassist_pv() from the R package findr, and function cit.cp() from the R package cit on each set to test whether there is a causal model as in V_1_→T_1_→T_2_ or V_1_→T_2_→T_1_ (**Supplementary Table 5**).

The Genotype-Tissue Expression (GTEx) Project was supported by the Common Fund of the Office of the Director of the National Institutes of Health, and by NCI, NHGRI, NHLBI, NIDA, NIMH, and NINDS. The gene expression data used for the analyses described here were obtained from the GTEx Portal (https://www.gtexportal.org/home/datasets; GTEx_Analysis_2016-01-15_v7_RNASeQCv1.1.8_gene_tpm.gct.gz) on 10/24/2017, the genotype data were available through dbGaP accession number phs000424.v7.

### A.10 Properties of MRPC

A causal graph with a mixture of directed and undirected edges is essentially an equivalent class of directed acyclic graphs (DAGs) that have the same likelihood. However, the search problem of learning the DAG with the highest likelihood when the number of parent nodes is greater than 1 has been proven to be NP-complete^51^, the hardest computational problem. Learning even just the equivalent classes of a DAG with the number of parent nodes being greater than 1 is also NP-complete^52^, as the space of equivalent classes of DAGs is super-exponential^49^ in the number of nodes. Therefore, the PC algorithm and similar algorithms get around the computational issue with local searches. Although it is not known theoretically that these PC algorithms achieve the global optimality defined by, for example, the likelihood, it has been shown that the PC algorithm is consistent^49^: with a large sample size, the PC algorithm is expected to recover the true graph. In particular, consistency of the PC algorithm is essentially consistency of the step of graph skeleton inference, as this step contains all the statistical inference^53^. Since MRPC uses essentially the same procedure for skeleton inference as the PC algorithm, MRPC is also consistent.

### A.11 Multiple genetic variants of the same phenotype

MRPC currently does not directly deal with multiple genetic variants associated with the same molecular phenotype. For network inference, we recommend using the variant with the strongest association, or merging the multiple variants to create a haplotype variant with the haplotypes being the new genotypes (e.g., two SNPs in linkage disequilibrium, each having three genotypes, can be merged into one variant with genotypes 00, 01, 02, 10, 11, 12, 20, 21, and 22).

### A.12 MRPC and other PMR-based methods

Although our MRPC employs the PMR, it is fundamentally different from other PMR-based methods. Most of the methods incorporating the PMR fall into two classes. One class, including cit and findr, is called *mediation-based methods* that require individual-level data, generally do not estimate the causal effect sizes, and can infer networks of multiple phenotypes (e.g., a network of gene expression). The other class of methods are called *MR methods^22^* that can be applied to individual-level data as well as summary statistics, estimate the causal effect sizes, and generally focus on three-node graphs with one node being the genetic variant, and the other two nodes being phenotypes of interest. Both classes of methods employ the PMR and focus on the “causal model”, in which exposure acts as the mediator. Although our MRPC method is closer to the mediation-based methods according to the characteristics described above, the notion of “mediation” is less relevant to our method; only Model 1 considers the “causal model”, and therefore one of the two genes acts as the mediator (**Fig. 1a**). More importantly, with our method we consider the PMR as a way to define plausible causal relationships and to put constraints on the space of possible graphs. As a result, our method can recover a variety of causal relationships, instead of the few that other PMR-based methods can identify (**Fig. 2b**).

### Code availability

MRPC is implemented in an R package (v1.0.0) at https://cran.r-project.org/web/packages/MRPC/index.html.

## ACKNOWLEDGMENTS

We thank Jonathan Pritchard, Anand Bhaskar, Towfique Raj, Boxiang Liu, Deigo Calderon, and Xiaoyue Wang for helpful discussions, Boxiang Liu for processing the GTEx genotype data, and Diego Calderon for detailed and constructive comments on an earlier version of the paper. We also thank Lingfei Wang for help with the findr package, Huange Wang for providing the QPSO code.

